# Neurons in the mouse deep superior colliculus encode orientation/direction through suppression and extract selective visual features

**DOI:** 10.1101/092981

**Authors:** Shinya Ito, David A. Feldheim, Alan M. Litke

## Abstract

The superior colliculus (SC) is an integrative sensorimotor structure that contributes to multiple vision-dependent behaviors. It is a laminated structure; the superficial SC layers (sSC) contain cells that respond to visual stimuli, while the deep SC layers (dSC) contain cells that also respond to auditory and somatosensory stimuli. Despite the increasing interest in mice for visual system study, the differences in the visual response properties between the sSC and the dSC are largely unknown. Here we used a large-scale silicon probe recording system to examine the visual response properties of neurons within the SC of head-fixed, awake and behaving mice. We find that both the sSC and dSC cells respond to visual stimuli, but dSC cells have three key differences. (1) The majority of the dSC orientation/direction selective (OS/DS) cells have their firing rate *suppressed* by drifting sinusoidal gratings (negative OS/DS cells) rather than being stimulated like the sSC cells (positive OS/DS cells). (2) Almost all the dSC cells have complex-cell-like spatial summation nonlinearity, and a significantly smaller fraction of the positive OS/DS cells in the dSC respond to flashing spots than those in the sSC. (3) The dSC cells lack Y-like spatial summation nonlinearity unlike the sSC cells. These results provide the first description of cells that are suppressed by a visual stimulus with a specific orientation or direction, show that neurons in the dSC have properties analogous to cortical complex cells, and show the presence of Y-like nonlinearity in the sSC but their absence in the dSC.

**Significance statement:** The superior colliculus receives visual input from the retina in its superficial layers (sSC) and induces eye/head orientating movements and innate defensive responses in its deeper layers (dSC). Despite their importance, very little is known about the visual response properties of dSC neurons. Using highdensity electrode recordings and novel model-based analysis, we find that the dSC contains cells with a novel property; they are suppressed by the orientation or direction of specific stimuli. We also show that dSC cells have properties similar to cortical “complex” cells. Conversely, cells with Y-like nonlinear spatial summation properties are located only in the sSC. These findings contribute to our understanding of how the SC processes visual inputs, a critical step in comprehending visually-guided behaviors.

**Acknowledgements:** This work was supported by the Brain Research Seed Funding provided by UCSC and from the National Institutes of Health Grant NEI R21EYO26758 to D. A. F. and A. M. L. We thank Michael Stryker for training on the electrophysiology experiments and his very helpful comments on the manuscript, Sotiris Masmanidis for providing us with the silicon probes, Forest Martinez-McKinney and Serguei Kachiguin for their technical contributions to the silicon probe system, Jeremiah Tsyporin for taking an image of neural tissues and the training of mice, Jena Yamada, Anahit Hovhannisyan, Corinne Beier, and Sydney Weiser, for their helpful comments on the manuscript.

## Introduction

The superior colliculus (SC) is a midbrain structure that integrates inputs from different sensory modalities and controls multiple vision-dependent behaviors. In addition to its well-known role in eye/head movements (Sparks, 1986), the mouse SC contributes to other visually-evoked behaviors such as escape and/or freezing in response to a looming object (Shang et al., 2015; Wei et al., 2015) and quick suspension of locomotion (Liang et al., 2015). In fact, mice that develop without a cortex can perform visually-guided memory tasks, suggesting that the SC has functions that are often attributed to the cortex (Shanks et al., 2016).

The SC is organized into several synaptic laminae, each of which has distinct inputs and outputs (May, 2005). The stratum opticum (SO) contains retinal and cortical axons and roughly divides the SC into two parts: the SO and above are the superficial SC (sSC), which receives inputs from the retina and the primary visual cortex (V1) and responds to visual stimuli; and below the SO is the deep SC (dSC), which contains multimodal cells that respond to visual, auditory and/or somatosensory stimuli. The response properties of the mouse sSC cells have been well characterized and show selectivity to stimulus features such as orientation or direction (Wang et al., 2010; Inayat et al., 2015). Moreover, (Gale and Murphy, 2014) identified four cell types in the sSC that differ in their response properties and axonal targets, suggesting that different visual features are segregated into different cell types.

Compared to the sSC, much less is known about the visual response properties of the dSC cells. In mice, the only electrophysiological studies of the dSC are the original studies conducted by (Drager and Hubel, 1975a, 1975b, 1976) and (Drager, 1975), which reported that dSC cells have large receptive fields, are more frequently DS, and can be multimodal. These early results provide important hints of the dSC cell properties, but the number of recorded neurons is small (< 100) and the OS/DS properties were poorly quantified. In addition, there are other response properties that give clues to a cell's function that have not been explored in the dSC. These include two distinct types of nonlinear spatial summation properties: the cortical complex-cell-like nonlinearity measured by drifting sinusoidal gratings (C-like nonlinearity, Skottun et al., 1991) and the retinal Y-cell-like nonlinearity measured by contrast reversing gratings (Y-like nonlinearity, Hochstein and Shapley, 1976).

To fill this knowledge gap, we have used large-scale silicon probe neural recordings to examine the visual response properties of the SC cells of head-fixed, awake, behaving mice watching a variety of visual stimuli. We also developed a model-based analysis that provides a thorough parameterization and simple significance tests of the OS/DS response properties. Using a dataset with 1445 identified neurons from both the sSC and the dSC, we discovered several novel features of the mouse SC. First, we found that OS and DS cells exist in both the sSC and the dSC. However, the majority of the OS/DS cells in the dSC have their spiking activity *suppressed* by drifting sinusoidal gratings that drift in their preferred orientation/direction (negative OS/DS cells). Negative OS/DS cells have not been previously reported in the mammalian visual system. Second, we found that a larger fraction of dSC cells have C-like nonlinearity than sSC cells, and that approximately one-third of dSC positive OS/DS cells lack any response to a flashing spot, unlike most sSC positive OS/DS cells. These differences between sSC and dSC cells are analogous to those between simple and complex cells in the visual cortex. Finally, we found cells with Y-like nonlinearity in the sSC, but not in the dSC. Taken together, our large-scale recordings provide an important step to gain a comprehensive analysis of the visual response properties of cells in the mouse SC with the goal of understanding how this area contributes to visually-guided behaviors.

## Materials and Methods

All procedures were performed in accordance with the University of California, Santa Cruz (UCSC) Institutional Animal Care and Use Committee. We recorded the activity of the collicular neurons of head-fixed, awake, behaving mice using 128-electrode or 256-electrode multi-shank silicon probes. Our experimental procedures were previously described (Shanks et al., 2016) and are detailed below.

*Animal preparation and experimental setup*. 2–5 months old C57BL/6 male mice were used in this study. Before the recording date, we implanted a stainless steel head plate on the mouse's skull; this allowed us to fix the mouse's head to the recording rig. Mice were trained to behave freely on a spherical floating ball treadmill (Dombeck et al., 2007; Niell and Stryker, 2010) for at least 4 training sessions (30 min per training session, each session on a different day). On the day of the recording, the mouse was anesthetized with isoflurane (3% induction, 1.5–2% maintenance), and a craniotomy (~1.5 mm diameter) was performed in the left hemisphere at a site that was 0.6 mm lateral from the midline and 3.7 mm posterior from the bregma. During the surgery, the mouse eyes were covered with artificial tears ointment (Rugby), which was removed before recovery from anesthesia with a wet cotton swab. The incision was covered with 2% low melting point agarose in saline to keep the exposed brain tissue from drying. The probe was inserted through the cortex toward the SC; the end of the probe was located 1800–2300 μm below the cortical surface. Recordings were started 30 minutes after probe insertion. During the recording sessions, the mouse was allowed to behave freely on the treadmill. We recorded only once from each mouse.

A visual stimulus monitor (Samsung S29E790C; 67.3 cm × 23.4 cm active display size; mean luminance: 32 cd/cm^2^; refresh rate: 60 Hz; gamma corrected) was placed 25 cm away from the right side of the mouse (Fig. 1A). Activity in the SC was identified by displaying either an ON (white) or OFF (black) flashing spot on the monitor and determining whether visual responses were localized to a restricted area within the visual field. The monitor position and angles were adjusted so that the receptive field was at the center of the monitor and the monitor plane was perpendicular to the line of sight.

**Figure 1:**
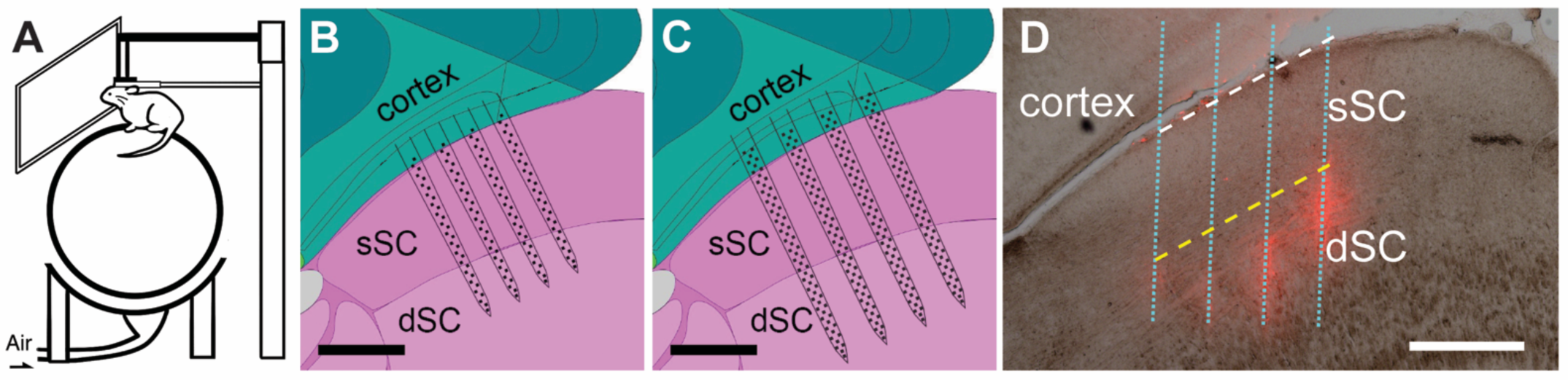
Experimental setup. A: A drawing of the recording setup. During the recording, the headfixed mouse is awake, behaving on a floating spherical treadmill and watching a variety of visual stimuli. B: A schematic of a 128-electrode, 4-shank silicon probe (150 μm shank pitch, each shank is 65 μm wide and 23 μm thick; 2 columns of electrodes on each shank, 20 μm pitch; 16 electrodes on each column, 50 μm pitch; 775 μm active length, 470 μm lateral extent; used for 5 of 20 recordings). The background image is a sagittal section of the SC, taken from the Allen Mouse Brain Atlas (Lein et al., 2007) with labels modified to match the terms in the present study. The scale bar is 400 μm. C: A schematic of a 256-electrode, 4-shank silicon probe (200 μm shank pitch, each shank is 86 μm wide and 23 μm thick; 3 columns of electrodes on each shank, 20 μm pitch; 21–22 electrodes on each column, 50 μm pitch; 1050 μm active length, 640 μm lateral extent; used for 15 of 20 recordings). The scale bar is 400 μm. D: A photograph of a parasagittal section of the SC after recording, indicating the probe location as marked by DiI. The dotted cyan lines indicate the recording shank locations reconstructed by the DiI traces (red channel). A dashed white line indicates the electrophysiologically estimated surface of the SC, which agrees well with the actual surface. A dashed yellow line indicates the estimated border between the sSC and the dSC, which is 400 μm below the estimated surface of the SC. The scale bar is 400 μm.

*Silicon probe electrophysiology*. The silicon probe recording was performed as described in our previous report (Shanks et al., 2016) with an important modification. Instead of 64 and 128-electrode 2-shank silicon probes, we used 4-shank probes with 128 electrodes and 256 electrodes (Fig. 1B,C). All the probes used for our recordings were kindly provided by Prof. Masmanidis at UCLA (Du et al., 2011). A small amount of DiI was put on the back of the probe shanks. The location of the shanks were histologically reconstructed after recordings using the DiI traces (Fig. 1D). A ground wire was placed on the skull near the craniotomy site by submerging it into the agarose. The voltage traces from all the electrodes were amplified and sampled at 20 kHz using an RHD2000 256-channel recording and data acquisition system (Intan Technologies). The visual stimuli were synchronized to the recorded neuronal activity via electrical pulses sent from the visual stimulation computer to the data acquisition board.

*Visual stimuli*. Four different visual stimuli were used to evaluate the visual responses of the SC neurons: (1) individual 10° diameter flashing circular spots on a 10 × 7 grid with 10° spacing. The 500 ms flashes were either ON (white) or OFF (black) on a gray background at mean luminance and a 500 ms gray screen was inserted after each stimulus presentation. The stimulus contrast and the location on the grid were chosen in a random order. Each pattern (every combination of the grid locations and the contrasts) was repeated 12 times (Wang et al., 2010); (2) drifting sinusoidal gratings: the parameters were the same as those used in (Niell and Stryker, 2008). The sinusoidal gratings were moving in 12 different directions (30° spacing) with 6 spatial frequencies in a range, 0.01–0.32 cycles/degree (cpd), with geometric steps. The temporal frequency and duration were 2 Hz and 1.5 s, respectively. We also presented full field flickering (0 cpd) and gray screen. Each pattern was presented in a random order and repeated 12 times; (3) contrast-reversing sinusoidal gratings: the gratings had 7 spatial frequencies in a range, 0.02–1.28 cpd, with geometric steps. The amplitudes were sinusoidally modulated at 4 Hz. Two different spatial phases (0° and 90°), and 4 different orientations (0°, 45°, 90°, and 135°) were used for each spatial frequency. Each pattern was presented in a random order and repeated 10 times; and (4) a contrast-modulated noise movie adapted from (Niell and Stryker, 2008): Briefly, a Gaussian white noise movie was generated in the Fourier domain with the spatial frequency spectrum set to *A*(*f*)= 1/(*f+f_c_*), with *f_c_* = 0.05 cpd, and a sharp low-pass cutoff at 0.12 cpd. The temporal spectrum was flat with a sharp low-pass cutoff at 4 Hz. This frequency domain movie was then transformed into the temporal domain and a 0.1 Hz sinusoidal contrast modulation was added. The movie was originally generated at 128 × 64 pixels, and expanded to the size of the stimulus monitor with smooth interpolations between pixels.

In addition to these stimuli, a contrast-alternating checkerboard stimulus (0.04 cpd square wave, alternating at 0.5 Hz) was employed to collect the local field potentials (LFPs), which were used to estimate the surface location of the SC (see the ‘Electrophysiological identification of the SC surface’ section).

*Spike-sorting and extraction of the local field potential*. For spike-sorting and local field potential analysis, we used custom-designed software as described previously (Litke et al., 2004;Shanks et al., 2016). A level 5 discrete wavelet filter (Wiltschko et al., 2008) (cutoff frequency ~313 Hz) was applied to the recorded data. The high-pass part was used for single unit identification after motion-artifact removal, whereas the low-pass part was used for the LFP analysis. The average motion artifact shape was estimated as a function of time by averaging the signals of all the recording channels. The estimated artifact was then subtracted from each channel with a multiplicative factor that minimized the root mean square of the channel.

Individual neuron identification was based on a previously developed method (Litke et al.,2004). The waveform of a spike detected on a “seed” electrode was combined with the time-correlated waveforms on the neighboring electrodes. Principal Component Analysis was used to extract the most significant variables for spike sorting from the waveform measurements. These variables were then clustered using the expectation maximization algorithm. After the first fit, the number of clusters was reduced one by one and refit until it no longer lowered the Bayesian information criterion (Fraley and Raftery, 1998). To remove duplicates and bad clusters, we used the contamination index (> 0.3, Litke et al., 2004), isolation distance (< 20, Schmitzer-Torbert et al., 2005), L-ratio (> 0.1, Schmitzer-Torbert et al., 2005), spike-correlation (> 0.25, Litke et al., 2004), similarity of electrophysiological images (>0.95 inner product), and the firing rate (requires > 0.1 Hz in the first and last 5-minute segments of the visual stimulus). When responses to multiple visual stimuli are compared, we required the firing rate criterion to be satisfied for all the visual stimuli. Once the neurons were identified, the positions of the neurons were estimated by fitting a 2-dimensional circular Gaussian to the spatial extent of the average spike amplitudes (i.e. electrophysiological images (EIs); see (Litke et al., 2004) for details) of the neurons.

*Electrophysiological identification of the SC surface*. We identified the surface of the SC using the LFPs recorded on each electrode. A flashing checkerboard stimulus generates a strong LFP in the area where the sSC is located (Zhao et al., 2014). We used the following steps to identify the surface line of the SC. (1) On a vertically aligned column of electrodes, average the evoked LFPs induced by the checkerboard contrast reversal to get the LFP as a function of the depth and the time after the reversal (Fig. 2A). (2) Choose the time when the LFP has the maximum negative amplitude and, at this specific time, obtain the LFP amplitude as a function of depth (Fig. 2B). (3) Choose the depth where the LFP is half maximum and closer to the top of the shank. This point is defined as the surface position for the column. (4) Repeat (1) through (3) for all available electrode columns (8 columns for 128-electrode probes; 12 columns for 256-electrode probes), and obtain a surface point for each column. (5) Fit a line to the identified surface points using the method of least squares. This electrophysiologically estimated SC surface agrees well with the actual surface of the SC (Fig. 1D, dashed white line). The perpendicular distance from this line defines the depth of the cells in the SC.

**Figure 2:**
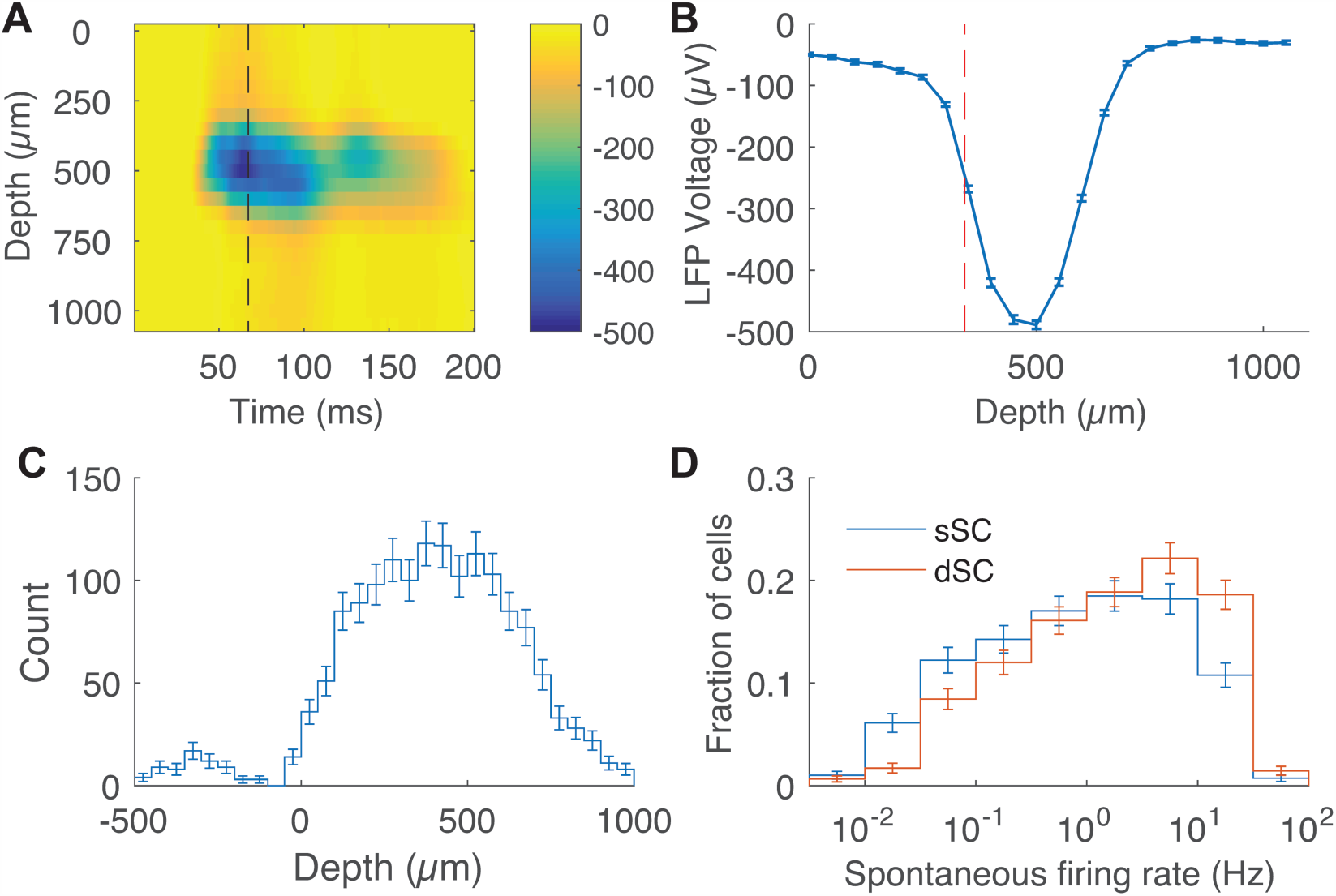
SC surface estimation, and firing rate distribution. A: An image of the average LFP evoked by a checkerboard stimulus. The image represents data collected from electrodes in one 22-electrode column. The color indicates the LFP voltage. The vertical axis gives the depth from the top electrode. The horizontal axis gives the time after the contrast reversal of the checkerboard stimulus. The black dashed line indicates the time when the amplitude of the LFP response is maximized. B: The LFP voltage as a function of depth at the time indicated by the black dashed line in (A). The red dashed line indicates the estimated surface of the SC for the electrode column, where the LFP voltage is 1/2 of its maximum amplitude. Error bars are SEM over 600 trials. C: Number of identified neurons as a function of the depth. The neurons with a negative depth are cortical neurons. The gap between the cortex and the SC right above zero confirms that the depth of the SC surface is estimated well. D: Distribution of the spontaneous firing rate of the sSC (blue) and dSC (red) cells. The dSC cells are more spontaneously active.

This procedure will produce a similar result to that of the current source density (CSD) analysis. The CSD analysis is an established method for identifying physiological landmarks of anatomical structures (Niell and Stryker, 2008; Zhao et al., 2014). The CSD is the second order derivative of the LFP with respect to depth. Another method to identify the surface of the SC is to use a zero-crossing point of the CSD. However, if you estimate the LFP amplitude as a Gaussian function of depth (with mean μ and standard deviation σ), which is a reasonable model given the shape shown in Fig. 2B, the zero-crossing point of the CSD, μ ± σ (with σ ~ 100 μm), is not very different from the half-maximum point, μ ± 1.18 σ. In addition, the half-maximum point has the following advantages: (1) resilience to the asymmetry of the LFP shape; (2) robustness to the outlier amplitudes outside the region of interest; and (3) methodological/computational simplicity. Therefore, we decided to use the half-maximum point method instead of the CSD method.

We defined the border between the sSC and dSC as 400 μm in depth measured perpendicularly from the surface. This is based on the change of the physiological properties in our own data, as well as the fact that a border line between the SO and the stratum griseum intermedium (SGI) was drawn consistently at ~400 μm in other studies (Phongphanphanee et al., 2008; Hong et al., 2011; Zhao et al., 2014). We do not distinguish any further sublamina of the SC—our dSC includes areas that are sometimes referred to as the intermediate and/or deep SC.

*Spontaneous firing rate*. We define the spontaneous firing rate of an individual neuron as the firing rate while the mouse watches a gray screen. An accurate measurement of the spontaneous firing rate is important because the significance and sign of the neuronal response is evaluated relative to this rate. We tried two different methods for evaluating the spontaneous firing rate. Our drifting grating stimulus consists of 1.5 s stimulus periods and 0.5 s intervals. In the 12 stimulus periods, we displayed a blank gray screen pattern to evaluate the spontaneous firing rates (1.5 s × 12 = 18 s total time for evaluation). In addition, we also evaluated the spontaneous firing rates during the intervals after removing the first 0.2 s, which is affected by switching from the stimulus to the gray screen (0.3 s × 887 = 266.1 s). The two spontaneous firing rates did not differ significantly (p > 0.01) for most neurons (92%). Therefore, we used the spontaneous firing rates evaluated by the intervals as they are more precise.

*Modeling of the orientation/direction selectivity with a χ^2^ fit*. We used *χ*^2^ minimization to fit our model functions to the firing rate of a cell to stimuli with different directions (Direction tuning curve; DTC). The *χ*^2^ is defined as,

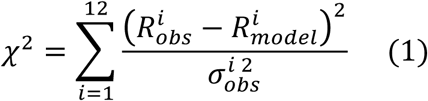

where the sum is over all the twelve directions i, R^i^_obs_ is the observed average firing rate, R^i^_model_ is the estimated firing rate from the model functions that are defined below, and σ^i^_obs_ is the SEM of the observed firing rate. The preferred spatial frequency of a neuron was chosen as the frequency that caused the most significant firing rate change from the spontaneous rate, either positive or negative.

The Python SciPy ‘curve_fit’ function was used for fitting functions. (In order to have proper error propagations, the version needs to be ≥ 0.17.) A goodness-of-fit test was done for each fit with the evaluation of the p-value. The p-value distribution is a useful indicator of how well the model fits the data (Fig. 3E).

**Figure 3:**
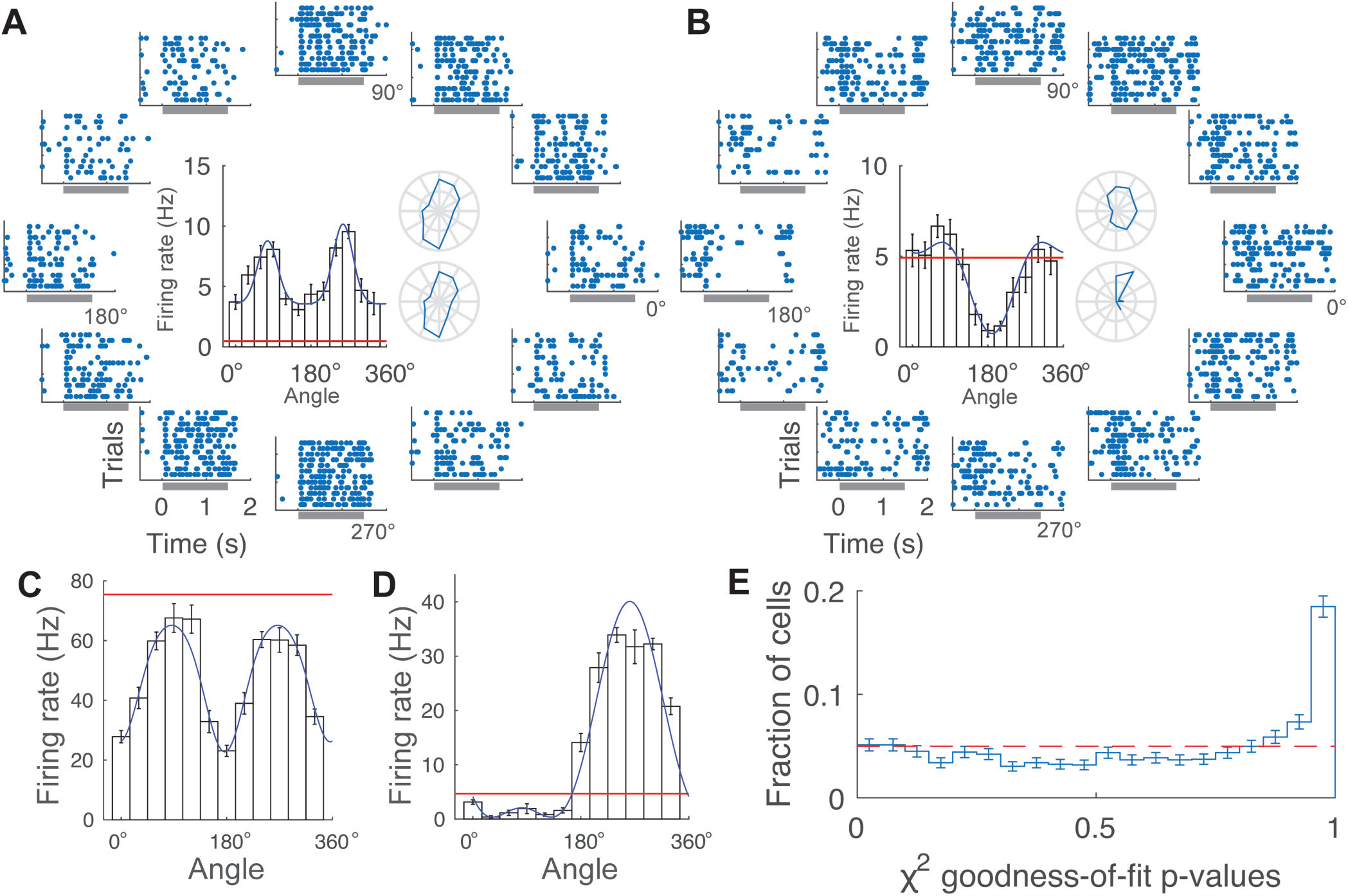
Model fit examples. A: Example drifting gratings responses of a positive OS cell. The circularly placed raster plots are the responses to the gratings moving toward the corresponding directions. The gray horizontal bars indicate the beginning and end of the stimulus. The inset bar graph indicates the average response firing rate for each direction (black histogram), the spontaneous firing rate (red line), and the result of a function fit (blue solid line). The error bars indicate the SEM of 12 trials. Two polar plots to the right of the histogram indicate the polar plot representation of the response histogram. The polar plots are the direction tuning curves (DTCs) of the cell without (top) and with (bottom) the spontaneous firing rate subtracted. B: The same set of figures as A for a negative DS cell. The firing rate is significantly lower than the spontaneous rate around 190°. Note that the polar plots no longer represent the correct characterization of the response property of this neuron. C: A response histogram of a negative OS cell. Note that the spontaneous firing rate is above the responses for all the directions, thus this neuron would be neglected with the traditional approach of spontaneous firing rate subtraction. D: A response histogram of a DS cell that responds both positively and negatively relative to the spontaneous firing rate. This neuron also has a particularly large range of response firing rate, and the firing rate seems to saturate near its preferred direction. Such a response causes deformation in the sinusoidal or Gaussian shapes, and results in a poor fit (p = 1 − 2 × 10^−6^). E: Distribution of the fit p-values (of *χ^2^*) for all the neurons. If the fit is consistent with the data points and their corresponding errors, the expected distribution is flat. We can estimate the fraction of neurons that were modeled poorly by the fit by counting the number of neurons above the expected flat line. ~17% of the neurons exceeded the flat line, indicating that ~83% of the neurons were well-modeled by our fit models.

Two model functions were used for the fit.

<u>Wrapped Gaussians:</u>

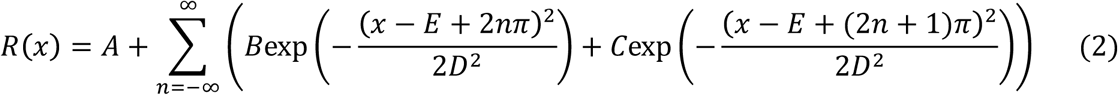

where R(x) is a periodic function: R(x+2π) = R(x). In the actual implementation, the range of n is set to −3 to 3, which serves as a practical approximation of this function for 0 < x < 2 pi.

As previously reported, the Gaussian fit does not always converge if the parameters are unbounded (Mazurek et al., 2014). We introduced fit parameter boundaries that are similar to (Mazurek et al., 2014).

1. 0 < A < max(DTC) (To avoid blowup of the baseline, which happens when the width is large)
2. (bin width) / 2 < D < π / 2 (min: To avoid overfitting by shrinking Gaussians max: To avoid excessive overlapping of the adjacent Gaussians.)
3. -4π < E < 4π (To avoid E getting out of the defined function)

<u>Sinusoid:</u>

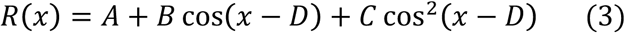

There are no parameter restrictions for the sinusoidal model.

The fit parameters were evaluated with an error matrix (Hessian matrix). As previously noted (Mazurek et al., 2014), the error is not trustworthy when the fit parameter is at the manually set boundaries; however, even if some parameters are at the boundaries, the errors of the other parameters are still valid. We used the error values only when the fit parameters are not at their boundaries.

In order to compare the results of the fits from these two different fit functions, we calculated various OS/DS properties from the fit parameters (Table 1). When arithmetic calculations were performed on the parameters, the errors were appropriately propagated using both the variance and the covariance of the parameters. A cell with a significant (positive or negative) DS amplitude (p < 0.001) was classified as a DS cell, and a non-DS cell with a significant OS amplitude (p < 0.001) was classified as an OS cell. We used a significance threshold at p = 0.001 in order to reduce the fraction of false positive OS/DS cells in the subsequent analysis. For example, assuming that 20% of all the cells are OS cells, applying p = 0.01 and p = 0.001 thresholds will result in having ~4% and ~0.4% of the candidate OS cells, respectively, to be false-positives.

**Table 1:**
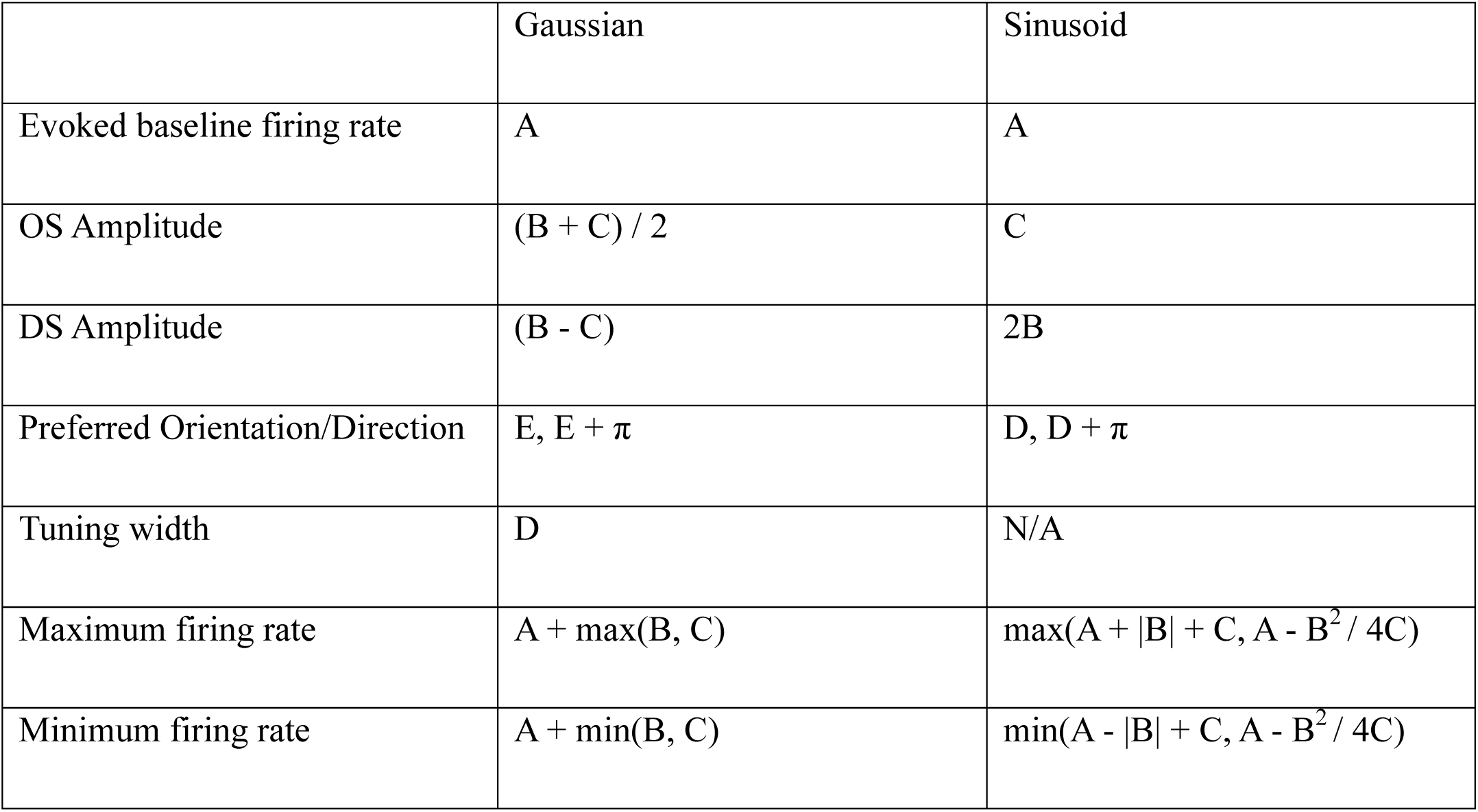
Calculation of the OS/DS properties by fit parameters.

The maximum and minimum firing rates determine whether a neuron had a positive response, a negative response or both. If the maximum/minimum firing rate is significantly higher/lower (p < 0.01) than the spontaneous firing rate, the neuron has a positive/negative response, respectively.

This model-based method resolves the issues of the traditional methods used by others, which subtract the spontaneous firing rate from the response (Niell and Stryker, 2008; Wang et al., 2010; Inayat et al., 2015). The treatment of the negative firing rate caused by this subtraction is not explicitly explained in these studies. If the negative part is truncated, it loses information of the negative part; if the negative part is left as negative, the definition of the preferred angle—the weighted sum of the phase—becomes ill-defined. Also, the resulting parameters (orientation/direction selectivity index; OSI/DSI) were not checked for significance. (Mazurek et al., 2014) introduced a method of significance tests, but the relation with the spontaneous firing rate was not discussed in their study. Our model-based method provides a proper treatment of both the spontaneous firing rate and the significance tests.

*Goodness of fit and goodness of the model*. The goodness of fit was evaluated for each fit. If the fit is good, the minimized *χ*^2^ values should follow the *χ*^2^ distribution. The two models have a different number of parameters (5 for the Gaussian fit and 4 for the sinusoidal fit) and therefore a different number of degrees of freedom (7 and 8, respectively). In order to cross-compare the *χ*^2^ distributions with different degrees of freedom, we calculated the goodness of fit p-values using the cumulative probability distribution function of the *χ*^2^ distribution. Low p-values indicate overfitting or overestimated errors; high p-values indicate too few parameters, an incorrect model or under-estimated errors.

*Analysis of flashing spot response*. A flashing spot can elicit spikes for most of the sSC cells (Wang et al., 2010). We used a previously described method for the flashing spot response analysis (Shanks et al., 2016). In this study the significance of a response was evaluated at each flashing spot grid location. The poststimulus time histogram was calculated using a 50 ms bin size. The response was considered significant if the firing rate associated with a bin exceeded the mean firing rate of the neuron during the entire flashing spot stimulus by at least 5σ. Here σ is the SD of the firing rate based on Poisson statistics. There are 70 grid locations, 2 contrasts and 10 time bins. After Bonferroni correction, it corresponds to a significance level of p~0.0004.

We characterized only the positive response to the flashing spots. With the binning size (50 ms) and the number of trials (12) that we used in our experiments, a negative response could only be a statistically significant amplitude for neurons with a very high spontaneous firing rate (≥ 42 Hz), which most neurons did not have.

*Analysis of contrast reversing gratings*. The contrast reversing gratings is a spatial sinusoidal pattern that changes contrast sinusoidally over time. By using a wide range of spatial frequencies, including frequencies that exceed the cells’ receptive field spatial resolution limit, it is possible to characterize the Y-like nonlinear spatial summation property (Hochstein and Shapley, 1976; Petrusca et al., 2007). Responses at the 4 Hz stimulus temporal frequency (F_1_) and the second harmonic frequency (F_2_) were characterized as a function of spatial frequency using a method previously described (Hochstein and Shapley, 1976; Petrusca et al., 2007). Namely, for each spatial frequency, the phase dependence was taken into account by taking the maximum F_1_ value, and the mean F_2_ value, over the two spatial phases.

Neurons are considered to have Y-like nonlinearity of spatial summation when they have a significant response in the F_1_ component at the lowest spatial frequency (0.02 cpd, p < 0.01), and have a significantly stronger F_2_ response in at least one of the higher frequencies (p < 0.01, with Bonferroni correction). In a primate retina study, the nonlinearity index (maximum value of F_2_/F_1_ ratio over the spatial frequencies) has been used (Petrusca et al., 2007). The nonlinearity index is a good indicator of the Y-like cells in the retina, where most of the neurons are either ON cells or OFF cells. In the SC, many of the cells are ON-OFF cells (Wang et al., 2010), which have a strong F_2_ component even in a low spatial frequency range. The method used in the present study ensures that a cell has proper characteristics of a Y-like cell—a strong F_1_ component at a low spatial frequency, and a strong F_2_ component at a high spatial frequency.

*Analysis of the contrast modulated noise movie*. The contrast modulated noise movie is an effective stimulus to find out how cells respond to contrast. In order to characterize the contrast response of the cells, we used a function fit method similar to the analysis of the orientation selectivity. We considered the zero-contrast timings as the starting times of the stimulus, and constructed a poststimulus time histogram (PSTH) for each neuron. The same sinusoidal function (equation 3), with a 10 s temporal period, was fit to the PSTH (*χ*^2^ fit using SEM of the PSTH) to get the amplitude of the linear response (model parameter B). Among the cells with a significant linear response, those with a maximum firing rate at 4–6 s (when the contrast is maximum) are considered as stimulated-by-contrast cells; those with a maximum firing rate at 0–1 s and 9–10 s are considered as suppressed-by-contrast cells. ‘Other’ cells include those with a non-significant linear component, and those with a maximum firing rate timing that does not meet the above criteria.

*Blind analysis*. Blind analysis is an effective method for reducing the false-positive reporting of results (Klein and Roodman, 2005; MacCoun and Perlmutter, 2015). (Please note that the analysis is “blind”, not the mice!) We looked for specific and/or significant features in a randomly subsampled set of 10 out of the 20 recordings. After specifying the features for further investigation, and freezing the individual analyses, we unblinded the other 10 blinded recordings to check if the significance of the results was reproduced. All the results shown in this paper passed the significance test before and after unblinding. We report the results of the combined data.

## Results

### Large scale silicon probe recordings create a large dataset of visually responsive neurons throughout the depth of the SC

We recorded from the SC of awake, behaving mice on a spherical treadmill (Fig. 1A) using large-scale silicon probes (Fig. 1B,C). The active length of the probe is long enough to record from both the sSC and the dSC simultaneously. The surface location of the SC was estimated by the LFPs induced by the alternating checkerboard stimulus (Fig. 2A,B; see ‘Electrophysiological identification of the SC surface’ section in Materials and Methods), and the neurons were divided into the sSC (depth < 400 μm) and the dSC (depth ≥ 400 μm) neurons. The high spatial density, 128- and 256-electrode silicon probe recordings identified a total of 1445 SC neurons (687 superficial, 758 deep) in 20 mice, with an average number of 0.43 ± 0.03 isolated single neurons per electrode in the SC. Fig. 2C shows the distribution of cells throughout the depth of the SC. The distribution of the spontaneous firing rate is shown in Fig. 2D; the mean and the SEM in the log space were 10^−010^ ^± 003^ and 10^019^ ^± 003^ Hz (~0.8 and ~1.6 Hz) in the sSC and the dSC, respectively (8 sSC cells with zero spontaneous spikes were excluded from the calculation). The dSC cells had significantly higher spontaneous firing rates than the sSC cells (p = 7.2 × 10^−10^). Some analyses require visual stimuli that were used in only a subset of experiments; in those cases, the number of neurons used for the analysis is indicated in the corresponding sections.

### Characterization of orientation and direction selective cells using a novel model-based analysis

In order to extract the OS/DS response properties in terms of model parameters, we applied a modelbased OS/DS analysis to the recorded neurons (see Methods). This approach allows us to evaluate how well the model fits the data. By applying a p < 0.001 significance threshold for the fitted OS or DS amplitudes, as defined in Table 1, we found 302 (173 superficial, 129 deep) OS cells and 201 (135 superficial, 66 deep) DS cells. Fig. 3A-D show examples of the raster plots and/or tuning curves of OS and DS cells, including: an OS cell with a positive firing rate change (Fig. 3A), a DS cell with a negative firing rate change (Fig. 3B), an OS cell with a negative firing rate change (Fig 3C), and a DS cell with both a positive and a negative firing rate change (Fig 3D). (All firing rate changes are evaluated relative to the spontaneous firing rate; see ‘Spontaneous firing rate’ in Materials and Methods). The preferred spatial frequency of the OS/DS cells was 0.01 × 2^2.25^ ^± 0.09^ (~0.048) cpd in the sSC and 0.01 × 2^2.47^ ^± 0.10^ (~0.056) cpd in the dSC, showing no significant difference (p = 0.10). The fraction of cells that are OS/DS as a function of depth is plotted in Fig. 4A, B. In the sSC 25.1 ± 1.7% of the cells are OS and 19.6 ± 1.5% are DS; in the dSC 17.0 ± 1.3% of the cells are OS and 8.7 ± 1.0% are DS. The area close to the surface (< 150 μm) is enriched with DS cells, showing qualitative consistency with (Inayat et al., 2015) but differing in the estimated abundance. Inayat et al. reported 74% of the cells in 0–50 μm depth are DS cells, while we find only 44 ± 8% of our cells are DS. This difference could be due to different methods for identifying the DS cells and/or our use of a relatively high level of the significance threshold (p < 0.001), compared to the DSI method with the significance unchecked.

**Figure 4:**
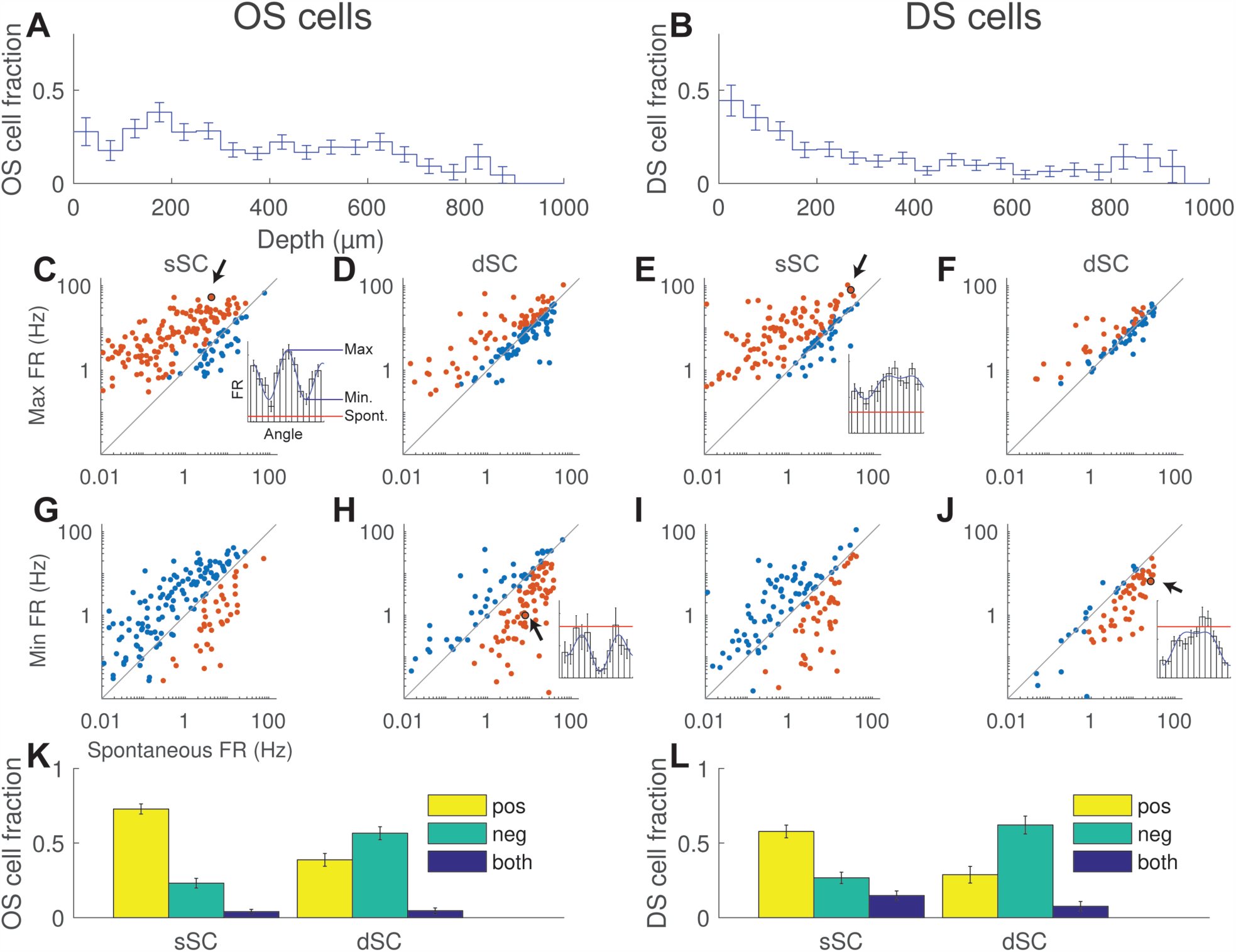
Response signs of the OS/DS cells. A, B: Fraction of cells that are OS (A) and DS (B) cells as a function of depth. The surface of the SC is enriched with DS cells. C, D: A scatter plot of the spontaneous firing rate (FR) vs. the maximum response to drifting gratings of OS cells in the superficial (C) and deep (D) SC areas. The red dots indicate neurons with a positive response (the maximum FR is significantly (p<0.01) above the spontaneous rate). A larger fraction of cells in the sSC have positive response. Inset shows the direction tuning curve of the cell that is arrowed in C. E, F: The same figures as C and D for DS cells. G, H: A scatter plot of the spontaneous FR vs. the minimum response to drifting gratings of OS cells in the superficial (G) and deep (H) areas. The red dots indicate neurons with negative response (the minimum firing rate is significantly (p<0.01) below the spontaneous rate). A larger fraction of cells in the dSC have negative response. I, J: The same figures as G and H for DS cells. K, L: Fractions of the cells that are OS (K) and DS (L) cells that have positive, negative, and both responses to the drifting gratings. The ‘positive’ OS/DS cells decrease, and the ‘negative’ OS/DS cells increase in the dSC (p < 0.001 for all combinations).

We determined the quality of the model fit for each individual cell by the *χ*^2^ per number of degrees of freedom (ndof) (with the expectation value *χ*^2^ = ndof and variance = 2 × ndof) and by the *χ*^2^ goodness-of-fit p-value. The overall quality of the model is evaluated by the distribution of the p-values for the set of fits for the population of neurons. If the model correctly represents the population data and the corresponding errors, a flat p-value distribution is expected as indicated in Fig. 3E. Using this rigorous method, we find that ~83% of the neurons are well-modeled and ~17% of the cells were poorly modeled (area above the red line in Fig. 3E).

### Cells that have negative orientation or direction selective responses are enriched in the dSC

Our model-based analysis also determines the sign (whether the firing rate change was positive, negative or both) of each cell's response to drifting grating stimuli. The model estimates the range of the firing rates in response to the drifting gratings. By comparing this range of the firing rate with the spontaneous firing rate, we determined whether the stimulus increased or decreased the firing rate of each cell (see Fig. 4C inset). Fig. 4C-J compare the minimum and maximum firing rates to the spontaneous firing rate of each OS and DS cell. The OS cells (C, D, G, H) and DS cells (E, F, I, J) show similar trends. Fig. 4C,E indicate that the sSC OS/DS cells respond to moving gratings with a positive change of the firing rate relative to their spontaneous rate, while Fig. 4H,J indicate that dSC OS/DS cells respond with a significant reduction of their firing rates. Fig. 4K, L summarize the results and show that, compared to the sSC, the fraction of the positive OS/DS cells decrease (OS: p = 4.4 × 10^−10^, DS: p = 3.5 × 10^−5^) and the fraction of the negative OS/DS cells increase in the dSC (OS: p = 6.3 × 10^−10^, DS: p = 5.5 × 10^−7^). Only a small fraction (7.6 ± 1.2%) of the OS/DS cells responded with both positive and negative firing rate change (the maximum firing rate was significantly above the spontaneous firing rate and the minimum firing rate was significantly below the spontaneous firing rate), while all the other cells had either positive or negative firing rate change.

### Suppressed-by-contrast cells are enriched in the dSC, and show little overlap with negative OS/DS cells

Suppressed-by-contrast (SuC) cells are a known example of a cell type with a decrease in firing rate in the presence of a high-contrast visual stimuli. As the negative OS/DS cells also have a firing rate reduction by the drifting gratings stimuli, an overlap between the SuC cells and the negative OS/DS cells might be expected. To measure the fractions of the stimulated-by-contrast (StC) cells and the SuC cells, we used the ‘contrast modulated noise movie’ adopted from (Niell and Stryker, 2008) (Fig. 5A). For this analysis, we used 551 neurons from 14 mice. The fraction of the StC and SuC cells as a function of depth is shown in Fig. 5B. The sSC consists mostly of StC cells (71 ± 3%) while the fraction of the SuC cells increases at ~400 μm in depth, and remains constant thereafter. The fractions of these cells in each category are summarized in Fig. 5C. In the dSC, the fraction of the StC cells decreases while the fraction of SuC cells and other cells increases.

**Figure 5:**
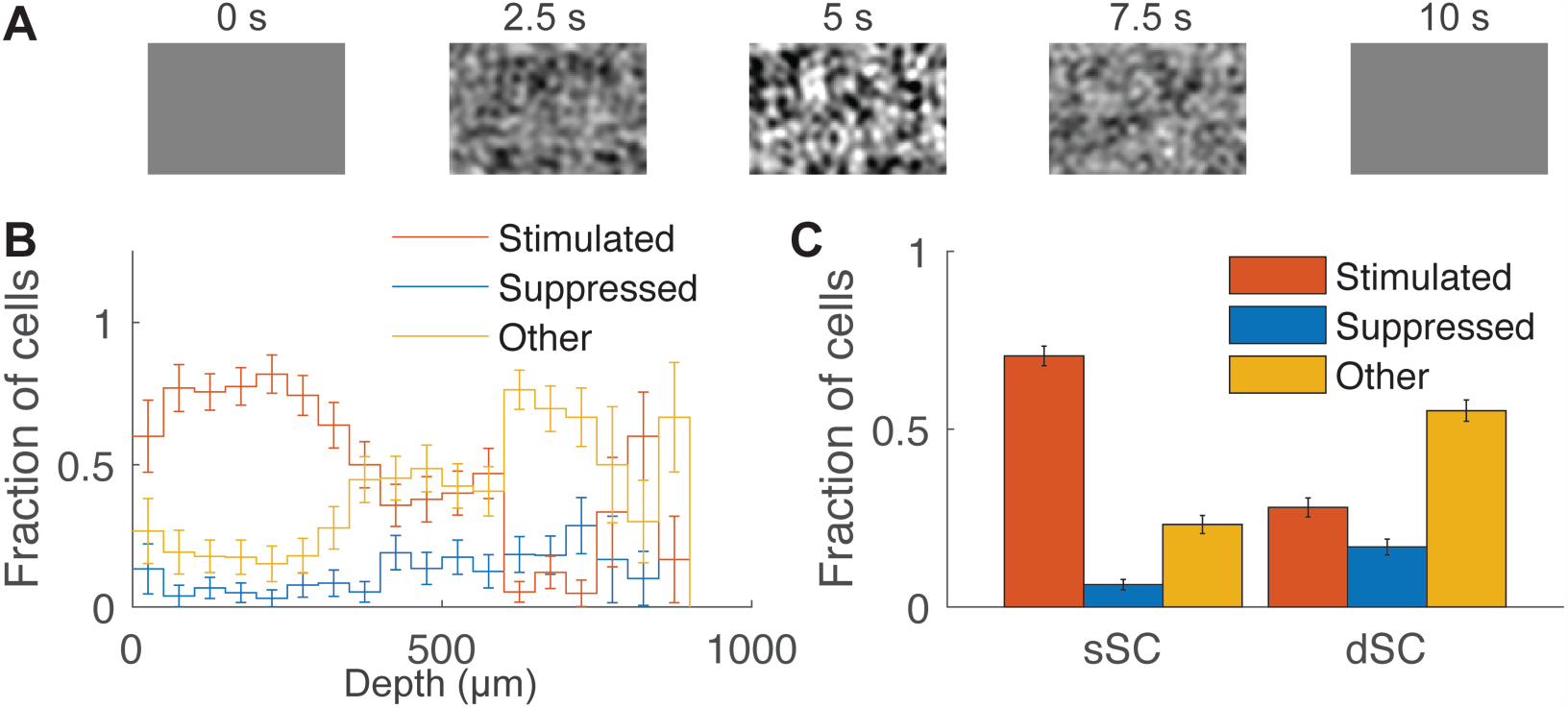
Suppressed-by-contrast cells. A: Panels showing the example stimulus pattern of the contrast modulated noise movie. The contrast of the movie is sinusoidally modulated over a 10 s period. B: Fraction of the cells, across the depth, that are conventional stimulated-by-contrast cells, suppressed-by-contrast cells, or cells in neither of these categories. The sSC is dominated by the stimulated-by-contrast cells. Soon after passing 400 μm, the fraction of stimulated-by-contrast cells decreases and the fraction of suppressed-by-contrast cells increases. C: Comparison of the fractions of the stimulated-by-contrast and suppressed-by-contrast cells and non-responsive cells in the superficial and deep areas. Note that the fraction of the suppressed-by-contrast cells and non-responsive cells increased in the deep area.

To determine if the negative OS/DS cells are a subset of the SuC cells, we measured the overlap between these cells. 77 ± 4% of the positive OS/DS cells were StC cells (chance value: 49 ± 2%), while only 22 ± 5% of the negative OS/DS cells were SuC (chance value: 12 ± 1%). Although they have more-than-chance overlap, a majority (78 ± 5%) of the negative OS/DS cells are not SuC cells. This result suggests that the negative OS/DS cells are not a subset of the SuC cells, and that they encode different specific visual features.

### Most of the dSC cells have complex-cell-like spatial summation nonlinearity

The ratio of the stimulus frequency response to the mean response, F_1_/F_0_, to a drifting sinusoidal grating, is a standard quantitative metric of response linearity and has been used to discriminate cortical simple cells from complex cells (Skottun et al., 1991). Here we define the cells with F_1_/F_0_ < 1 as C-like nonlinear cells and F_1_/F_0_ ≥ 1 as linear cells. This metric has been applied to mouse cortical cells (Niell and Stryker, 2008) and sSC cells (Wang et al., 2010). We have applied it to our data and found examples of both linear cells and C-like nonlinear cells in the SC (Fig. 6A). Note that the linear cell has a rhythmic response to the drifting gratings, while the C-like nonlinear cell elevates its firing rate regardless of the phase of the stimulus. Most cells with a linear response appear in the sSC (Fig. 6B) and most cells with a negative OS/DS response amplitude are C-like nonlinear (see Fig. 6C). Moreover, we see a clear difference between the cells with a positive response and the cells with a negative response (Fig. 6D). The dSC is enriched with negative OS/DS cells, which also means that they are enriched with C-like nonlinear cells. Even among the positive OS/DS cells, the distribution of the F_1_/F_0_ linearity was significantly different between the sSC and the dSC (p = 1.8 × 10^−6^, KS test; Fig. 6E), further highlighting the smaller fraction of linear cells and predominance of the C-like nonlinear cells in the dSC (Fig. 6F).

**Figure 6:**
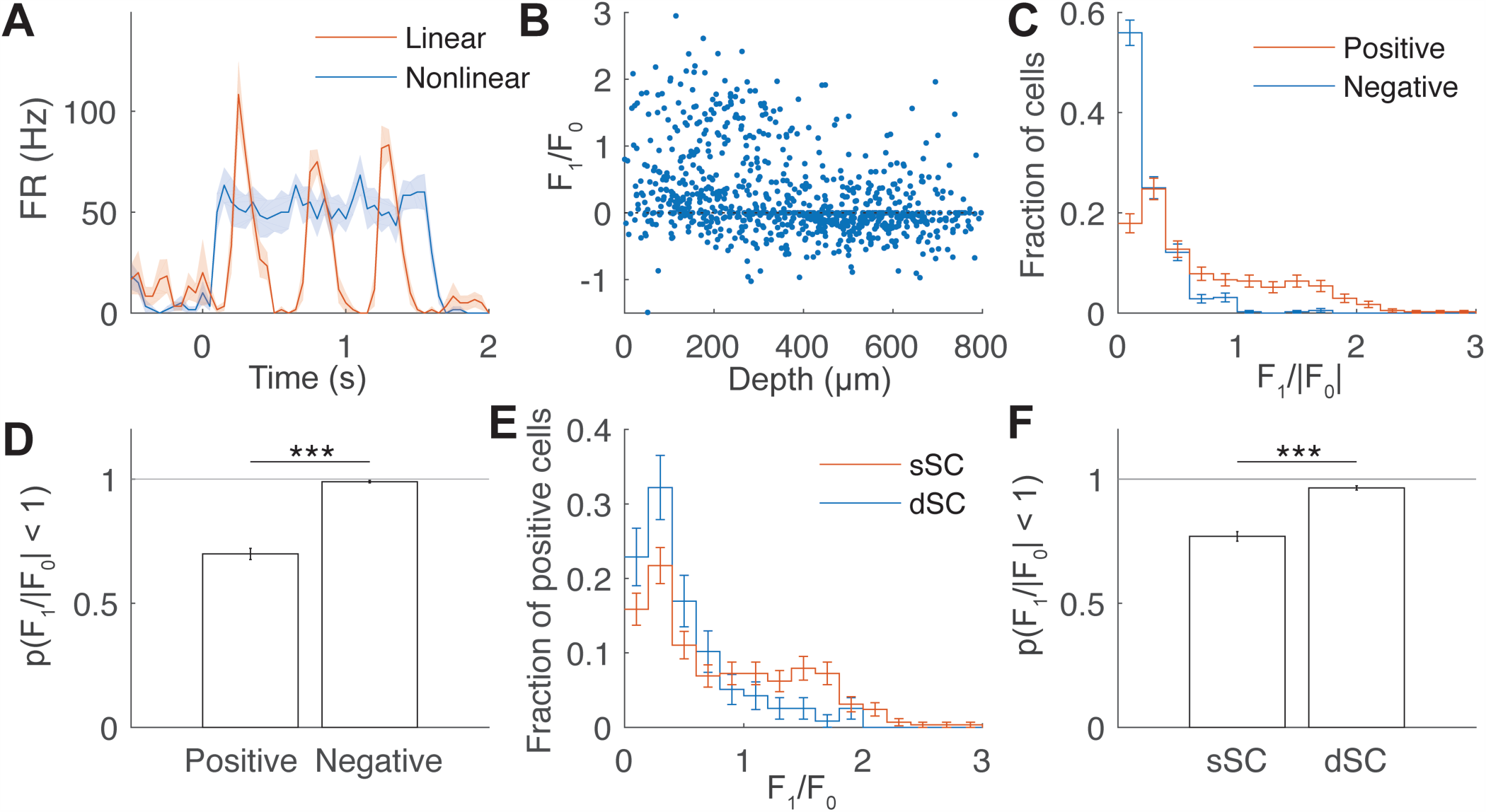
Difference of F_1_/F_0_ linearity between the sSC and the dSC. A: Typical temporal response of linear (red, F_1_/F_0_ = 1.97) and C-like nonlinear (blue, F_1_/F_0_ = 0.03) cells to drifting gratings. The linear cell responds rhythmically at the stimulus frequency (2 Hz), while the C-like nonlinear cell elevates the firing rate without a temporal pattern. B: A scatter plot of the depth and the F_1_/F_0_ nonlinearity. Note that F_0_ can have a negative value when the evoked firing rate is below the spontaneous firing rate, resulting in a negative F_1_/F_0_. C: Histograms of F_1_/F_0_ of neurons with positive and negative response. The distributions differ significantly (KS test, p=5 × 10^−30^). Most of the negative cells have a small value of F_1_/|F_0_|, indicating that they are C-like nonlinear. D: A bar graph comparing the fractions of the linear cells in positive and negative responding cells. The fraction of the cells that are linear is significantly larger among the cells with positive response than the cells with negative response (p = 7 × 10^−36^). E: Histograms of F_1_/F_0_ for positively responding cells. The distribution is different between the sSC and the dSC, showing a larger fraction of C-like nonlinear cells (F_1_/F_0_ < 1) in the dSC (KS test, p = 1.8 × 10^−6^). F: A bar graph comparing the fraction of the C-like nonlinear cells in the sSC and dSC. The dSC has a significantly larger fraction of the C-like nonlinear cells than the sSC (p = 1.9 × 10^−20^).

### Some dSC OS/DS cells do not respond to flashing spots

We also determined the response of the cells to a flashing spot stimulus (Fig. 7A, top), which is effective at stimulating sSC cells (Wang et al., 2010). For this analysis, we used a total number of 967 neurons. First, we quantified the fraction of the cells that are responsive to flashing spots in the sSC and the dSC. We found that the majority of the responsive cells respond both to the onset of a white flashing spot (ON stimulus) and the onset of a black flashing spot (OFF stimulus), consistent with a previous report (Wang et al., 2010). An example of an ON-OFF cell, responding to both white and black flashing spots, is shown in Fig. 7A (bottom). The percentages of the ON, OFF, ON-OFF and nonresponsive cells were 8 ± 1%, 16 ± 2%, 58 ± 2% and 17 ± 2% in the sSC and 5 ± 1%, 20 ± 2%, 31 ± 2%, and 44 ± 2% in the dSC, respectively. These non-responsive cells may respond to other visual stimuli, or they are non-visually-responsive cells such as auditory- and/or somatosensory-responsive cells.

**Figure 7:**
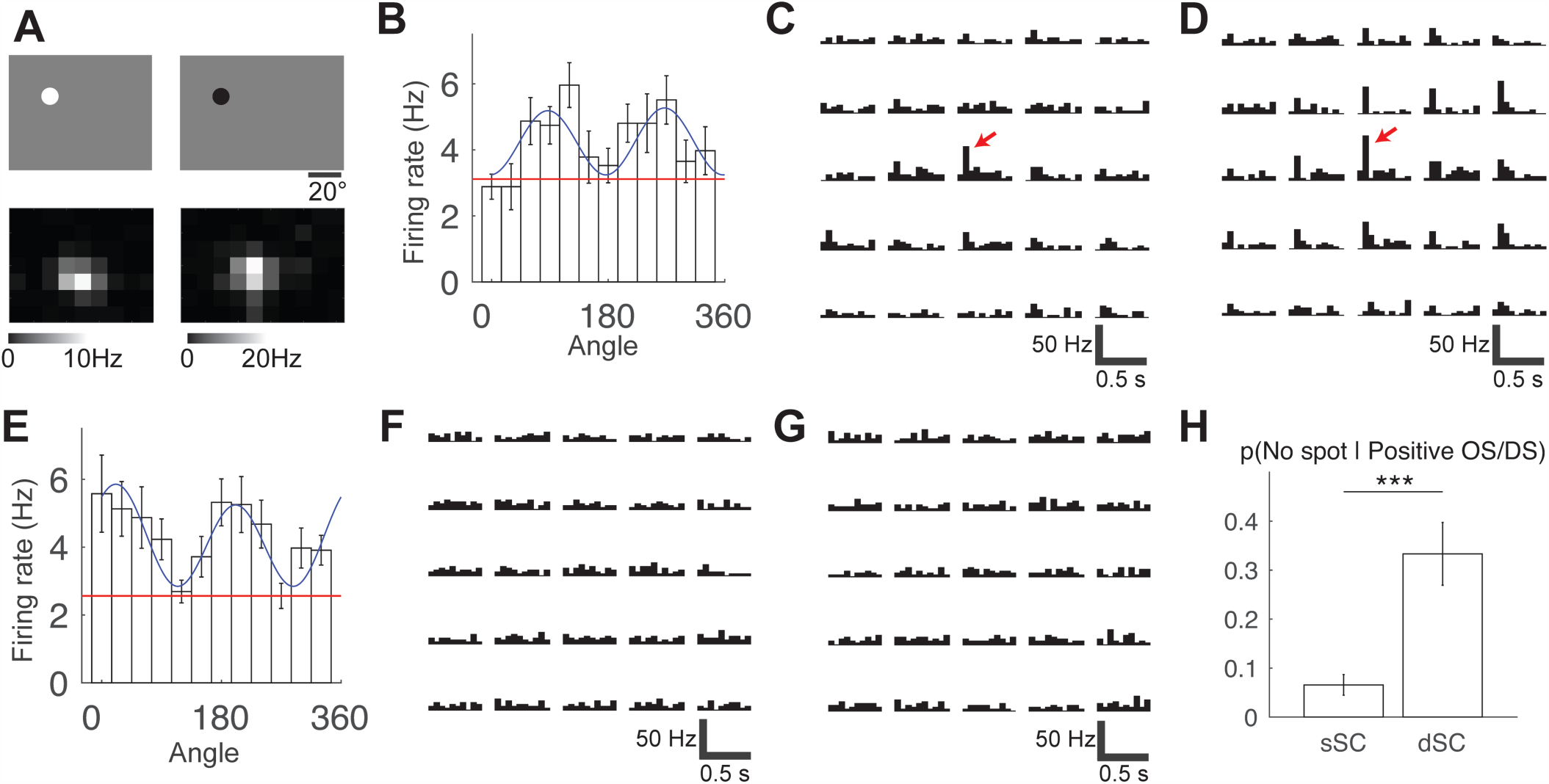
Lack of the flashing spot response in the dSC OS/DS cells. A: Flashing spot stimulus patterns and example responses. Example flashing white (top left) and black (top right) spot patterns are shown. These patterns were shown at different grid locations on the screen. Firing rates of an example neuron elicited by white (bottom left) and black (bottom right) spots at each grid location are shown in grayscale. This neuron responds to both black and white spots shown at a localized area of the screen. B: Response of an sSC OS cell to a drifting sinusoidal grating, and C, D: its response to a flashing white (C) or black (D) spot at different grid locations. A red arrow points to the crisp response of the cell to the appearance of a spot. The scale bars are 500 ms (horizontal) and 50 Hz (vertical). E, F, G: The same figures as B through D for a dSC OS cell. No response to either white or black flashing spots is observed. H: Fractions of the positive OS/DS cells that lack a flashing spot response in the sSC and dSC (***: p < 0.001). In the sSC, only 7 ± 2% of the positive OS/DS cells have no spot response, while in the dSC, 36 ± 7% of the positive OS/DS cells have no spot response.

Next, we characterized flashing spot responses of OS/DS cells, which are known to be visually responsive. For this analysis, we used only positive OS/DS cells, as it was statistically difficult to quantify negative responses to flashing spots (see ‘Analysis of flashing spot response’ in Methods) For this analysis, we used 209 positive OS/DS cells. We found that some of the dSC OS/DS cells do not respond to flashing spots. In order to illustrate the difference of the response between the sSC and the dSC, we chose an example cell from each region. Despite their similar orientation selectivity (Fig. 7B and E), these cells have very different responses to the flashing spot stimulus: the sSC neuron has a sharp response to the onset of the white and black flashing spots (Fig. 7C and D, respectively; red arrows), while the dSC cell has no response at all (Fig. 7F and G). The fraction of cells with no flashing spot response was 7 ± 2% and 33 ± 6% in the sSC and dSC, respectively (Fig. 7H). All the dSC OS/DS cells that lack a flashing spot response (n = 18) were also C-like nonlinear cells (defined above by F_1_/F_0_ < 1). To summarize, some C-like nonlinear cells in the dSC ‘selectively’ respond to gratings drifting in their preferred orientation/direction, but not to flashing spots.

### Y-like nonlinear cells are restricted to the sSC

The Y-like response, the nonlinear rectification of fine spatial texture, has not been previously characterized in the mouse brain. We characterized the Y-like nonlinearity and found a difference between the sSC and the dSC. For this analysis, we used 977 neurons. X-like cells show a response with the same frequency as the stimulus frequency at low spatial frequencies and this response decays as the spatial frequency increases (Fig. 8B,C). Y-like nonlinear cells show a frequency-doubling response at a high spatial frequency (Fig. 8A) as demonstrated in Fig. 8D, E. A significant fraction (8.3 ± 1.3%) of the cells in the sSC have this Y-like nonlinearity, while almost none of the cells (0.2 ± 0.2%) in the dSC have this property (Fig. 8F).

**Figure 8:**
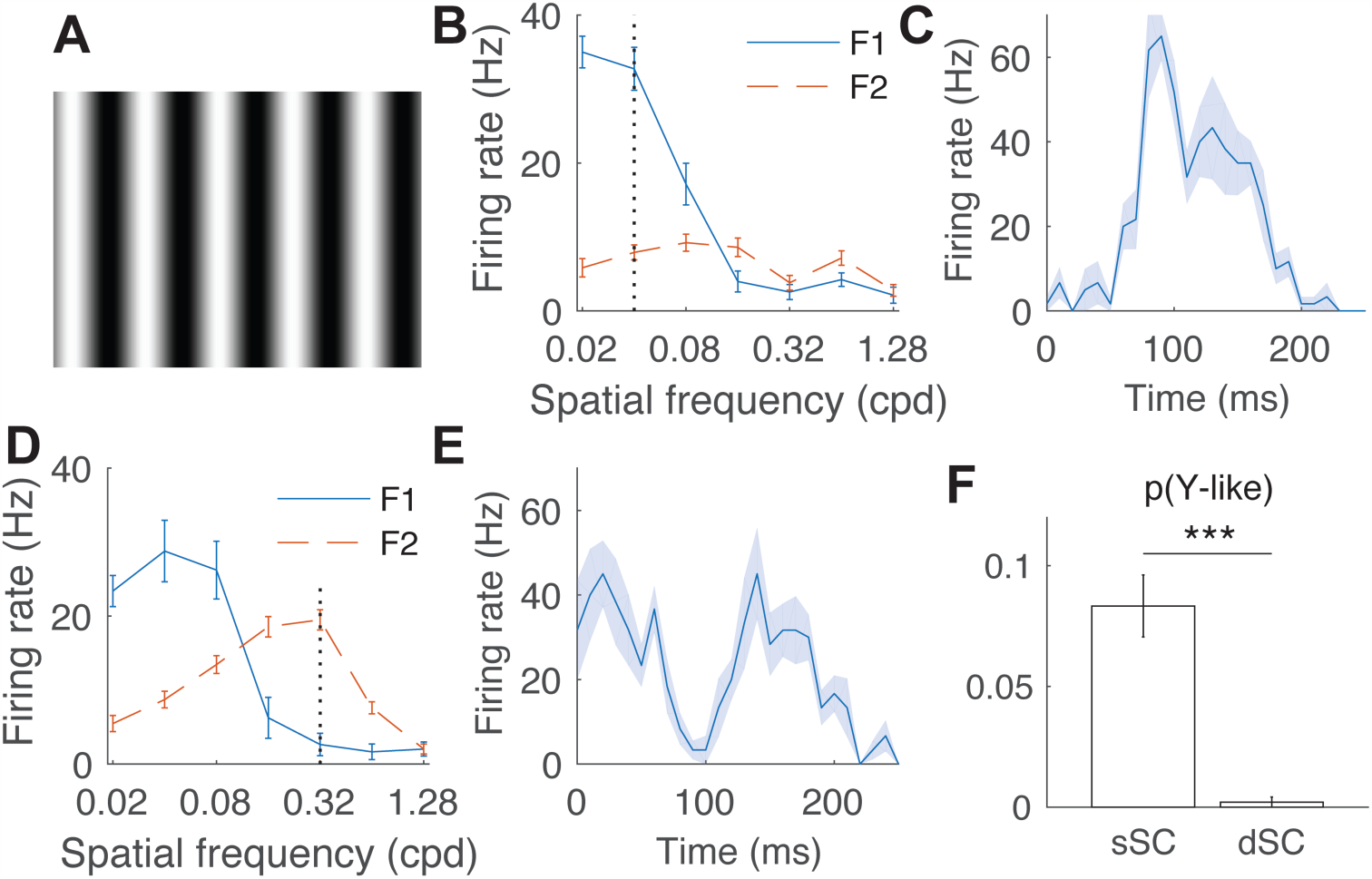
Y-like nonlinear cells appear only in the sSC. A: An example frame of a contrast reversing grating. Instead of moving across the screen, the contrast of the pattern changes sinusoidally at 4 Hz. B: The F1 (4 Hz) and F2 (8 Hz) Fourier components as a function of spatial frequency. This cell shows an X-like response, which has a strong F1 component. C: A time course of the response of the cell shown in (B). The response is taken at the spatial frequency indicated by the black dotted line in (B). In one stimulus cycle (250 ms), the cell shows only one peak. D: A figure corresponding to (B) for a Y-like cell. Although the F1 component is dominant at low frequency, the F2 component increases at high frequency and exceeds the F1 component. E: A time course of the response of the cell used for (D). Note that there are two peaks in one stimulus cycle. F: Fractions of the Y-like cells found in the sSC and the dSC. Almost all (42 of 43) of the Y-like cells were found in the sSC.

## Discussion

In this study, we used high-density silicon probe recordings in awake, behaving mice to examine the visual response properties of mouse SC cells, with particular attention to the less characterized dSC cells. In contrast to the sSC cells, the dSC cells have the following characteristics in visual response properties: (1) the majority of OS/DS cells encode their preferred orientations/directions by *suppression* of the firing rate; (2) almost all the cells have C-like nonlinearity and about one-third of the positive OS/DS cells lack a flashing spot response; and (3) Y-like nonlinearity is absent. These results provide important and novel details about the visual coding properties of the mouse SC as elaborated in the following discussion.

### Negative encoding in the dSC

Neurons in the early visual system (i.e. the retina, dorsal lateral geniculate nucleus (dLGN), V1, and sSC) typically encode information by a positive change of the firing rate. There are cases that nonpreferred stimuli lead to inhibition of these visual cells, but such inhibition is usually in conjunction with stronger excitation to the preferred stimuli. A known exception to such positive encoding is the ‘suppressed-by-contrast’ cells (also known as ‘uniformity detectors’) that reduce their firing rate with a high-contrast stimulus. In mice, they are known to exist in the retina (Tien et al., 2015), dLGN (Piscopo et al., 2013), and V1 (Niell and Stryker, 2010). We find these cells in the SC, at the level of 6.2 ± 1.5% in the sSC, increasing to 16 ± 2% in the dSC. One hypothesis is that these cells are reporting self-generated stimuli (saccade or blink) (Tien et al., 2015), but their detailed properties and functional roles remain unknown.

We found that 58 ± 4% of OS/DS cells in the dSC are ‘negative’ OS/DS cells, in contrast to the 25 ± 2% detected in the sSC. What is the role of these negative OS/DS cells? One hypothesis is that these are inhibitory cells that gate motor output. In fact, an in-vitro study showed that 74% of GABAergic neurons in the SGI (in our study called the dSC) have high spontaneous firing rates, and provide tonic inhibition in the local circuitry (Sooksawate et al., 2011). Unfortunately, inhibitory and excitatory neurons cannot be distinguished using the action potential waveforms in the SC, as was done in V1 (Niell and Stryker, 2008). Therefore, we will need further experiments to identify negative OS/DS cells as inhibitory or excitatory.

### Increased C-like nonlinearity and lack of response to flashing spots indicate that the mouse dSC cells are selective to visual features

Almost all of the cells (96.5 ± 0.9%) that respond to drifting gratings in the dSC have C-like nonlinearity; the fraction of the positive OS/DS cells that lack flashing spot responses was 7 ± 2% in the sSC and 33 ± 6% in the dSC. The dSC is characterized by predominance of C-like nonlinearity and lack of response to flashing spots compared to the sSC. Interestingly, these transitions are similar to those between the cortical simple and complex cells (nonlinearity: Skottun et al., 1991; lack of spot response: Hubel and Wiesel, 1962). Furthermore, both SC and V1 are laminar structures, and complex cells are enriched within their output layers (deep layers of the SC and layer 5 of V1, Niell and Stryker, 2008) This result suggests that the SC is performing visual feature extraction similar to those proposed for complex cells in V1.

In studies of the primate SC, few DS cells were found (Humphrey, 1968), and the visual response properties of the majority of the cells have been described as ‘event detectors’, which means that they respond to almost any stimuli in their receptive fields (non-selective to visual features) if the stimulus size is appropriate (Schiller and Koerner, 1971), and these properties do not change qualitatively in the dSC (Humphrey, 1968; Cynader and Berman, 1972; Goldberg and Wurtz, 1972). Such a visual response is not useful for finding out what the object is, but is useful for encoding the appearance and location of an object (Boehnke and Munoz, 2008), which are relevant information for producing an accurate gaze shift toward the object.

On the other hand, recent studies of anesthetized mice showed that the sSC cells are ‘selective’ to visual features (Wang et al., 2010; Gale and Murphy, 2014). We have confirmed visual selectivity in awake mice and showed that the dSC cells are even more selective than the sSC cells. Although we do not know why such a ‘selective’ versus ‘non-selective’ difference between mouse SC and primate SC arises, it is worth noting that there are two important differences between them. The primate SC is innervated by only ~10% of retinal ganglion cells (RGCs), while the mouse SC is innervated by at least 70% of RGCs (May, 2005), and the mouse does not have a fovea. It should also be noted that we did experiments with the visual cortex preserved. Therefore, some of the responses we find may be relayed from the visual cortex, as it has been shown that multiple areas of the visual cortex project to the SC (Wang and Burkhalter, 2013). Nevertheless, the mouse SC cells are more similar to the mouse or primate V1 cells than to the primate SC cells in the sense that they are selective to visual features, and this property may support the surprising visual task capability of mice that develop without a visual cortex (Shanks et al., 2016), and other visually guided behaviors (Liang et al., 2015; Shang et al., 2015; Wei et al., 2015).

### Cells with a Y-like nonlinear spatial summation property are found only in the sSC

We found cells with a Y-like nonlinearity in the sSC (8.3 ± 1.3%), but not in the dSC (0.2 ± 0.2%). This Y-like nonlinearity is a well-known property of the alpha/Y-type RGCs, originally found in the cat retina (Hochstein and Shapley, 1976), and subsequently in primates (de Monasterio, 1978; Petrusca et al., 2007), guinea pigs (Demb et al., 1999) and mice (Stone and Pinto, 1993). The Y-like nonlinearity of the sSC cells could be inherited from these RGCs. One of the alpha RGC types in mice, the transient OFF alpha RGCs (i.e. the PV-5 ganglion cells), have strong sensitivity to a looming spot stimulus, which imitates an approaching object (Münch et al., 2009). This stimulus causes an innate defensive response (Yilmaz and Meister, 2013) proposed to be through a retina-SC-lateral posterior nucleus (LP, also known as the pulvinar)-amygdala pathway (Shang et al., 2015; Wei et al., 2015). The SC cells that project to the LP have been identified as the wide-field cells, which do not project to the dSC (Gale and Murphy, 2014); this is consistent with our inability to detect dSC cells with these Y-like properties. Therefore, our results are consistent with the hypothesis that the wide-field cells in the sSC obtain their Y-like nonlinearity from the retina, and transmit them to the LP, but not to the dSC.

### Advantages of the model-based analysis and high-density silicon probes

We used simple model functions (see Methods) to describe the OS and DS cells. This model-based analysis extracts the response parameters of SC cells with their corresponding correlated uncertainties, and provides a method for significance tests between parameters. Using this analysis, we discovered a novel type of OS/DS cells that respond with suppression of their firing rate. This method can also be applied to the visual cortex, which may also contain negative OS/DS cells that have been overlooked.

In addition to this new method of analysis, the large-scale silicon probe recording system was important for the detailed characterization of the cells. For example, because the electrodes are dense (~32 μm between electrodes), the spike of a single cell is recorded simultaneously on several electrodes with different amplitudes and waveforms. These signals may seem redundant, but they improve both the single unit identification (Litke et al., 2004) and the cell position estimates. Furthermore, the largescale recording provides high statistical power. We recorded from 1445 cells, and this is the first electrophysiological study of the mouse SC conducted at this scale. This large dataset allowed us to characterize small populations such as negative OS/DS cells, suppressed-by-contrast cells and Y-like nonlinear cells, and to perform a blind analysis that increased confidence in the results.

## Conclusion

We characterized the visual response properties of both the sSC and the dSC using advanced hardware and software, and discovered that the dSC encodes stimulus orientation and direction with the suppression of the firing rate, performs cortex-like visual feature extraction, and lacks Y-like nonlinearity. This is the first report of the presence of negative OS/DS cells anywhere in the mammalian visual system, and they were discovered using the model-based analysis that we developed for the present study, combined with the large-scale silicon probe recording system. The cortex-like selective feature extraction suggests that the mouse SC contributes to visual functions that are often attributed to V1. These findings are an important step to understand how the SC processes visual images and contributes to visually-guided behavior of the mouse.

## References

Boehnke SE, Munoz DP (2008) On the importance of the transient visual response in the superior colliculus. Curr Opin Neurobiol 18:544–551.

Cynader M, Berman N (1972) Receptive-field organization of monkey superior colliculus. J Neurophysiol 35.

de Monasterio FM (1978) Properties of concentrically organized X and Y ganglion cells of macaque retina. J Neurophysiol 41.

Demb JB, Haarsma L, Freed MA, Sterling P (1999) Functional circuitry of the retinal ganglion cell's nonlinear receptive field. J Neurosci 19:9756–9767.

Dombeck DA, Khabbaz AN, Collman F, Adelman TL, Tank DW (2007) Imaging large-scale neural activity with cellular resolution in awake, mobile mice. Neuron 56:43–57.

Drager U, Hubel D (1975a) Responses to Visual Stimulation and Relationship Between Visual, Auditory, and Somatosensory Inputs in Mouse Superior Colliculus. J Neurophysiol:690–713.

Drager UC (1975) Receptive fields of single cells and topography in mouse visual cortex. J Comp Neurol 160:269–290.

Drager UC, Hubel DH (1975b) Physiology of visual cells in mouse superior colliculus and correlation with somatosensory and auditory input. Nature 253:203–204.

Drager UC, Hubel DH (1976) Topography of visual and somatosensory projections to mouse superior colliculus. J Neurophysiol 39:91–101.

Du J, Blanche TJ, Harrison RR, Lester HA, Masmanidis SC (2011) Multiplexed, high density electrophysiology with nanofabricated neural probes. PLoS One 6:e26204.

Fraley C, Raftery AE (1998) How Many Clusters? Which Clustering Method? Answers Via Model Based Cluster Analysis. Comput J 41:578–588.

Gale SD, Murphy GJ (2014) Distinct Representation and Distribution of Visual Information by Specific Cell Types in Mouse Superficial Superior Colliculus. J Neurosci 34:13458–13471.

Goldberg ME, Wurtz RH (1972) Activity of superior colliculus in behaving monkey. I. Visual receptive fields of single neurons. J Neurophysiol 35:542–559.

Hochstein S, Shapley RM (1976) Linear and nonlinear spatial subunits in Y cat retinal ganglion cells. J Physiol 262:265–284.

Hong YK, Kim I-J, Sanes JR (2011) Stereotyped axonal arbors of retinal ganglion cell subsets in the mouse superior colliculus. J Comp Neurol 519:1691–1711.

Hubel DH, Wiesel TN (1962) Receptive fields, binocular interaction and functional architecture in the cat's visual cortex. J Physiol 160:106–154.2.

Humphrey NK (1968) Responses to visual stimuli of units in the superior colliculus of rats and monkeys. Exp Neurol 20:312–340.

Inayat S, Barchini J, Chen H, Feng L, Liu X, Cang J (2015) Neurons in the Most Superficial Lamina of the Mouse Superior Colliculus Are Highly Selective for Stimulus Direction. J Neurosci 35:7992–8003.

Klein JR, Roodman A (2005) BLIND ANALYSIS IN NUCLEAR AND PARTICLE PHYSICS. Annu Rev Nucl Part Sci 55:141–163.

Lein ES et al. (2007) Genome-wide atlas of gene expression in the adult mouse brain. Nature 445:168–176.

Liang F, Xiong XR, Zingg B, Ji X ying, Zhang LI, Tao HW (2015) Sensory Cortical Control of a Visually Induced Arrest Behavior via Corticotectal Projections. Neuron 86:755–767.

Litke A, Bezayiff N, Chichilnisky EJ, Cunningham W, Dabrowski W, Grillo AA, Grivich M, Grybos P, Hottowy P, Kachiguine S, Kalmar RS, Mathieson K, Petrusca D, Rahman M, Sher A (2004) What does the eye tell the brain?: Development of a system for the large-scale recording of retinal output activity. IEEE Trans Nucl Sci 51:1434–1440.

MacCoun R, Perlmutter S (2015) Blind analysis: Hide results to seek the truth. Nature 526:187–189.

May PJ (2005) The mammalian superior colliculus: Laminar structure and connections. Prog Brain Res 151:321–378.

Mazurek M, Kager M, Van Hooser SD (2014) Robust quantification of orientation selectivity and direction selectivity. Front Neural Circuits 8:92.

Münch TA, da Silveira RA, Siegert S, Viney TJ, Awatramani GB, Roska B (2009) Approach sensitivity in the retina processed by a multifunctional neural circuit. Nat Neurosci 12:1308–1316.

Niell CM, Stryker MP (2008) Highly selective receptive fields in mouse visual cortex. J Neurosci 28:7520–7536.

Niell CM, Stryker MP (2010) Modulation of visual responses by behavioral state in mouse visual cortex. Neuron 65:472–479.

Petrusca D, Grivich MI, Sher A, Field GD, Gauthier JL, Greschner M, Shlens J, Chichilnisky EJ, Litke AM (2007) Identification and characterization of a Y-like primate retinal ganglion cell type. J Neurosci 27:11019–11027.

Phongphanphanee P, Kaneda K, Isa T (2008) Spatiotemporal profiles of field potentials in mouse superior colliculus analyzed by multichannel recording. J Neurosci 28:9309–9318.

Piscopo DM, El-Danaf RN, Huberman AD, Niell CM (2013) Diverse visual features encoded in mouse lateral geniculate nucleus. J Neurosci 33:4642–4656.

Schiller PH, Koerner F (1971) Discharge characteristics of single units in superior colliculus of the alert rhesus monkey. J Neurophysiol 34:920–936.

Schmitzer-Torbert N, Jackson J, Henze D, Harris K, Redish AD (2005) Quantitative measures of cluster quality for use in extracellular recordings. Neuroscience 131:1–11.

Shang C, Liu Z, Chen Z, Shi Y, Wang Q, Liu S, Li D, Cao P (2015) A parvalbumin-positive excitatory visual pathway to trigger fear responses in mice. Science (80-) 348:1472–1477.

Shanks JA, Ito S, Schaevitz L, Yamada J, Chen B, Litke A, Feldheim DA (2016) Corticothalamic Axons Are Essential for Retinal Ganglion Cell Axon Targeting to the Mouse Dorsal Lateral Geniculate Nucleus. J Neurosci 36:5252–5263.

Skottun BC, De Valois RL, Grosof DH, Movshon JA, Albrecht DG, Bonds AB (1991) Classifying simple and complex cells on the basis of response modulation. Vision Res 31:1078–1086.

Sooksawate T, Isa K, Behan M, Yanagawa Y, Isa T (2011) Organization of GABAergic inhibition in the motor output layer of the superior colliculus. Eur J Neurosci 33:421–432.

Sparks DL (1986) Translation of sensory signals into commands for control of saccadic eye movements: role of primate superior colliculus. Physiol Rev 66.

Stone C, Pinto LH (1993) Response properties of ganglion cells in the isolated mouse retina. Vis Neurosci 10:31–39.

Tien N-W, Pearson JT, Heller CR, Demas J, Kerschensteiner D (2015) Genetically Identified Suppressed-by-Contrast Retinal Ganglion Cells Reliably Signal Self-Generated Visual Stimuli. J Neurosci 35:10815–10820.

Wang L, Sarnaik R, Rangarajan K, Liu X, Cang J (2010) Visual receptive field properties of neurons in the superficial superior colliculus of the mouse. J Neurosci 30:16573–16584.

Wang Q, Burkhalter A (2013) Stream-Related Preferences of Inputs to the Superior Colliculus from Areas of Dorsal and Ventral Streams of Mouse Visual Cortex. J Neurosci 33:1696–1705.

Wei P, Liu N, Zhang Z, Liu X, Tang Y, He X, Wu B, Zhou Z, Liu Y, Li J, Zhang Y, Zhou X, Xu L, Chen L, Bi G, Hu X, Xu F, Wang L (2015) Processing of visually evoked innate fear by a non-canonical thalamic pathway. Nat Commun 6:6756.

Wiltschko AB, Gage GJ, Berke JD (2008) Wavelet filtering before spike detection preserves waveform shape and enhances single-unit discrimination. J Neurosci Methods 173:34–40.

Yilmaz M, Meister M (2013) Rapid innate defensive responses of mice to looming visual stimuli. Curr Biol 23:2011–2015.

Zhao X, Liu M, Cang J (2014) Visual Cortex Modulates the Magnitude but Not the Selectivity of Looming-Evoked Responses in the Superior Colliculus of Awake Mice. Neuron 84:202–213.

